# Network Topology Metrics Explaining Enrichment of Hybrid Epithelial Mesenchymal Phenotypes in Metastasis

**DOI:** 10.1101/2022.05.16.492000

**Authors:** Mubasher Rashid, Kishore Hari, John Thampi, Nived Krishnan Santhosh, Mohit Kumar Jolly

**Affiliations:** Center for BioSystems Science and Engineering, Indian Institute of Science, Bangalore 560012, India; Department of Mathematics, Indian Institute of Technology Kanpur, Kanpur, 208016, India; BS-MS Programme, Indian Institute of Science Education and Research, Pune, 411008, India; BS-MS Programme, Indian Institute of Science, Bangalore, 560012, India

**Keywords:** Metastasis, Epithelial Mesenchymal Transition, Hybrid Epithelial Mesenchymal Phenotype, Gene Regulatory Networks, Mathematical Modeling

## Abstract

Epithelial to mesenchymal transition (EMT) and its reverse mesenchymal to epithelial transition (MET) are hallmarks of metastasis. Cancer cells use this reversible cellular programming to switch among Epithelial (E), Mesenchymal (M), and hybrid Epithelial/Mesenchymal (hybrid E/M) state(s) and seed tumors at distant sites. Hybrid E/M cells are often more aggressive and metastatic than the “pure” E and M cells. Thus, identifying mechanisms to inhibit hybrid E/M cells can be promising in curtailing metastasis. While multiple gene regulatory networks (GRNs) based mathematical models for EMT/MET have been developed recently, identifying topological signatures enriching for hybrid E/M phenotypes remains to be done. Here, we investigate the dynamics of 13 different GRNs and identify which network topologies can enrich for hybrid E/M phenotype(s) across GRNs. We found that increasing negative feedback loops enhances the “hybridness” of these networks, but increasing the number of positive feedback loops or their specific combinations can decrease the frequency of hybrid E/M phenotype. Thus, our analysis identifies network topology based signatures that can give rise to, as well as prevent, the emergence of hybrid E/M phenotype in GRNs underlying EMP. Our results can have implications in terms of targeting specific interactions in GRNs as a potent way to restrict switching to the hybrid E/M phenotype(s) to curtail metastasis.

## Introduction

Metastasis – the spread of cancer cells from one organ to another – is a hallmark of cancer and causes over 90% of cancer-related deaths (Gupta and Massagué 2006). It begins with cancer cells being dislodged from the primary tumor, passing through the circulatory system and the lympho-vascular system, escaping the combat with immune system, and acclimatizing and proliferating at the secondary site (Massagué and Obenauf 2016). However, not all disseminated cancer cells contribute to metastasis; only <0.02% cancer cells can survive the chain of bottlenecks on the way to secondary tumor site and successfully metastasize (Luzzi et al. 1998; Cameron et al. 2000). Being a highly intricate process, metastasis remains poorly understood and is a key cause of therapy failure and the high fatality rate of solid tumors.

A key dynamical trait of metastasis is phenotypic plasticity – the ability of cancer cells to switch reversibly among different phenotypes (Jia et al. 2017a; Gupta et al. 2019). Epithelial Mesenchymal Plasticity (EMP) has been identified as an important branch of phenotypic plasticity, enriching metastasis through two programs: Epithelial Mesenchymal Transition (EMT) and its reverse, Mesenchymal Epithelial Transition (MET) (Bhatia et al. 2019; Tripathi et al. 2020c). Both these processes occur in normal developmental programs like embryogenesis, homoeostasis and wound healing (Nieto et al. 2016). EMT usually triggers a loss of cell-cell adhesion among epithelial (E) cells and concomitant gain of migratory and invasive traits associated with mesenchymal (M) state. Upon arrival at the secondary site, MET is considered to enable the disseminated M cancer cells to regain their epithelial traits and proliferate.

EMT was earlier thought to be a binary process, switching between epithelial and mesenchymal cells. However, recent studies have shown co-expression of epithelial and mesenchymal markers in individual cells in lung (Schliekelman et al. 2015; Jolly et al. 2016), colorectal (Hiew et al. 2018), renal (Sampson et al. 2014) and ovarian (Strauss et al. 2011), suggesting the existence of hybrid Epithelial/Mesenchymal (E/M) state (s). Hybrid E/M states have also been reported in primary tumors such as breast cancer (Yu et al. 2013), head and neck cancer (Puram et al. 2017), squamous cell cancer (Pastushenko et al. 2018), and colon cancer (Sacchetti et al. 2021) among others. Concomitantly, mathematical modeling of multiple gene regulatory networks (GRNs) for EMT/MET have endorsed that EMT proceeds via a cascade of hybrid E/M states, suggesting that not one but many hybrid E/M states can be acquired by cells rather stably along the EMT spectrum (Tian et al. 2013; Hong et al. 2015; Steinway et al. 2015; Font-Clos et al. 2018; Watanabe et al. 2019; Hari et al. 2020). Cancer cells in hybrid E/M state(s) are considered to be the “fittest” for the metastasis, given their association with tumor-initiating ability (stemness), resistance to multiple therapies, and association with worse patient survival (Biddle et al. 2016; Bierie et al. 2017; Jolly et al. 2018; Godin et al. 2020; Sahoo et al. 2021). Thus, identifying mechanisms that can restrict transition of E and M cells to hybrid E/M cells can be a crucial opportunity to design therapeutic targets to curtail metastasis.

Despite tremendous pathological implications, the hybrid E/M phenotype remains relatively poorly characterized. Recent efforts have identify some individual factors that can stabilize hybrid E/M phenotype(e) such as GRHL2, SLUG, NRF2, miR-129 and NFATc – referred to as the ‘phenotypic stability factors’ (PSFs) (Hong et al. 2015; Bocci et al. 2019; Silveira et al. 2020; Subbalakshmi et al. 2020, 2021; Norgard et al. 2021). However, these investigations focus on a handful of molecules at a time, thus overlooking how hybrid E/M phenotype can emerge as a function of network topology of the underlying GRN(s).

Here, we probe how network topology or design principles of GRNs impact the frequency of hybrid E/M phenotype(s). We investigate the dynamics of 13 different GRNs underlying EMT, varying in size and density from 4nodes/7edges (4N 7E) to 57 nodes/113 edges (57N 113E). Our discrete (parameter-independent) and continuous (parameter-agnostic) simulations suggest that rather than being the outcome of tuning the expression of a particular PSF, hybrid E/M phenotype can emerge due to collective interactions among the genes in these GRNs. Thus, we identified the network topology metrics of these GRNs that associate with abundance of hybrid E/M phenotype. Our network perturbation simulations revealed that increasing the negative feedback loops can significantly enhances the frequency of hybrid E/M phenotype, but increasing the number of both the positive feedback loops (PFLs) and *HiLoops* (Nordick and Hong 2021) – specific combinations of PFLs – decreases it. These results offer valuable insights to design therapies targeting interactions among the genes in EMT/MET networks, rather than modulating any particular gene of interest to curtail metastasis.

## Results

### “Hybridness” of EMP networks change with changing network-topology

To understand the emergence and frequency of hybrid phenotypes, we investigated 13 different GRNs underlying EMP and calculated the ‘hybridness’ of these networks. These networks can be categorized into 3 classes based on their size (number of nodes and edges). The first class is the four small networks (**Fig 1A i, Fig S1**). These networks were then combined to form networks of medium size (**Fig 1A ii, Fig S1**). In addition, we analysed five large-scale networks (**Fig 1A iii, Fig S1**). Each network is labelled with the number of nodes and edges in the network (ex: 4N 7E for the smallest network). We then simulated these networks and their single edge perturbations in which an edge is either deleted or its sign is reversed, i.e. an activatory (resp. inhibitory) edge is replaced by an inhibitory (resp. activatory) one (**Fig 1B**). These simulations are done using two complementary methods: a parameter agnostic, ordinary differential equation (ODE) based model – RACIPE (Huang et al. 2017), and a parameter-independent Boolean (logical) model using asynchronous update (Font-Clos et al. 2018).

**Figure 1.**
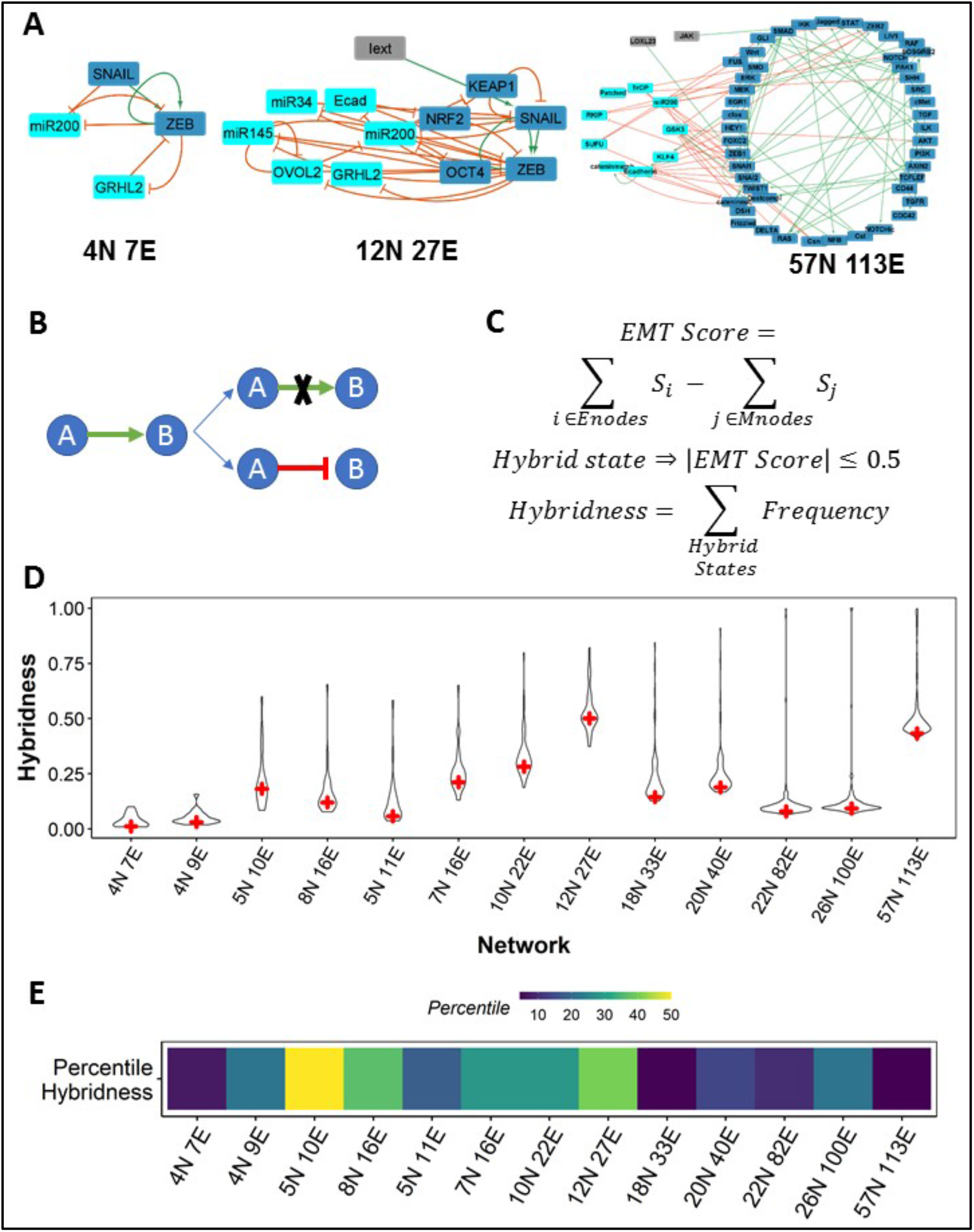
EMP networks show weak hybridness. **(A)** EMP network topologies used for analysis. **(B)** Depiction of two types of edge perturbations – edge deletion and edge-sign reversal. **(C)** Formula for the calculation of Hybridness for a network. Enodes and Mnodes are Epithelial and Mesenchymal nodes in the network respectively and S_i_ is the discretized expression level (0 or 1) of node i in state S. **(D)** Violin plots showing the distributions of hybridness obtained from RACIPE simulations for the ensemble of perturbed networks. ‘+’ sign indicates the hybridness of corresponding EMP (WT or unperturbed) network. **(E)** Percentile hybridness of WT EMP networks in the distributions shown in Fig 1D.

**Figure 1.**
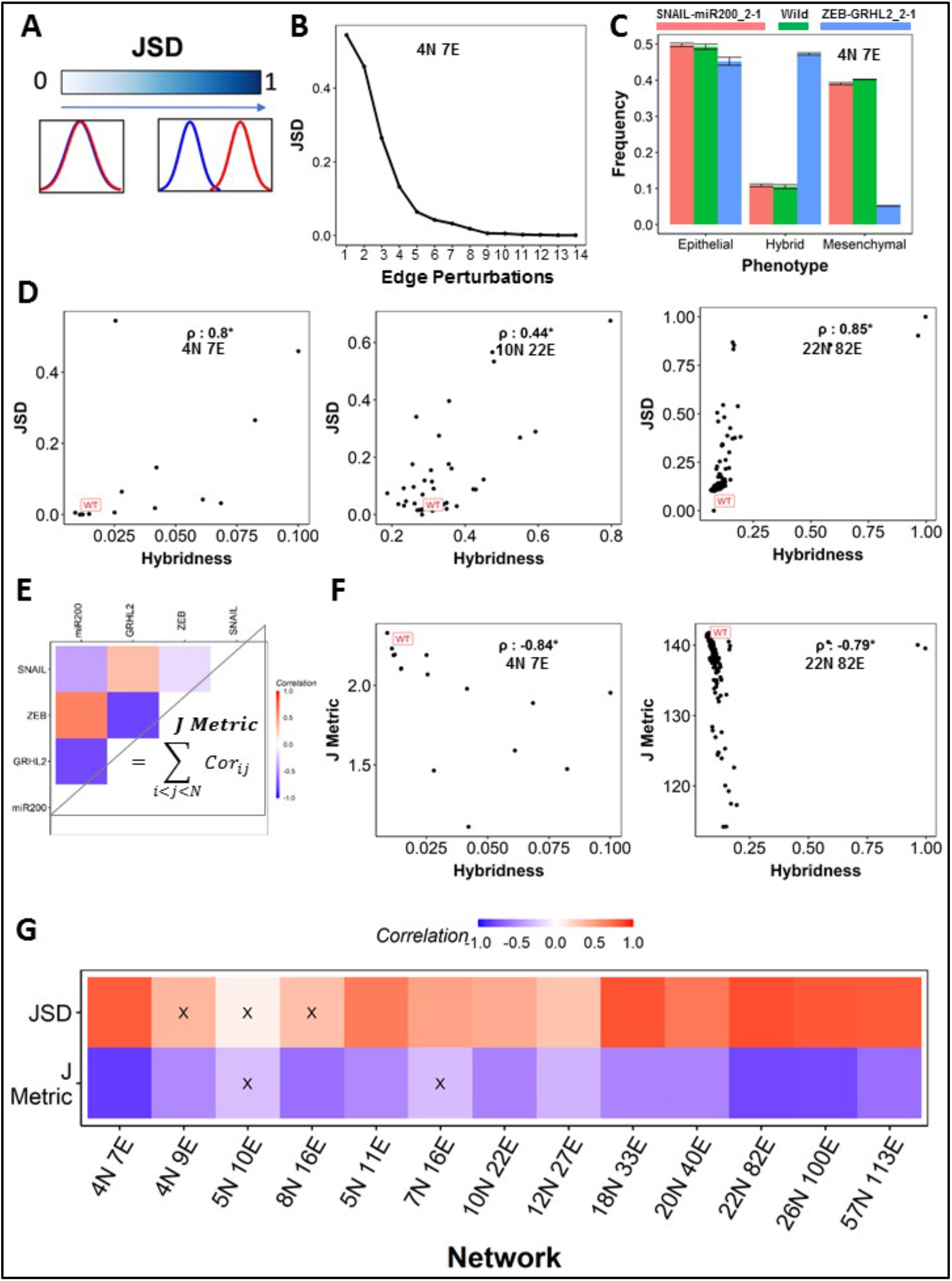
Change in global expression patterns correlates with change in hybridness. **(A)** Depiction of JSD as a measure of distance between two frequency distributions. **(B)** JSD values obtained for the single edge perturbation networks of 4N 7E EMP network. The terminology of network perturbation indices is given in Table S1. **(C)** Bar plot showing the frequency of Epithelial, Mesenchymal and Hybrid phenotypes for WT 4N 7E EMP network, perturbation with least JSD (Snail-miR200-2-1) and perturbation with highest JSD (ZEB-GRHL2-2-1). **(D)** Representative scatter plots between JSD and Hybridness for (i) 4N 7E, (ii) 10N 22E and (iii) 22N 82E. Each point corresponds to one perturbation. Spearman correlation coefficient is reported. **(E)** Depiction of J metric calculation with the example of 4N 7E network correlation matrix. **(F)** Representative scatter plots between J metric and Hybridness for (i) 4N 7E and (ii) 22N 82E. Spearman correlation coefficient is reported. **(G)** Spearman correlation between hybridness and JSD (top row) and J metric (bottom row). “X” represents p-value > 0.05. *: p-value < 0.05.

The ODE based simulations result in an array of continuous-valued steady states, which are then discretised based on their corresponding z-scores (the mean and standard deviation of expression values of nodes, calculated for ensemble of values across all the parameter sets). On the other hand, in Boolean (logical) framework, the value of a node can be either ON (1) or OFF (0). Boolean simulations are carried out using input edge threshold-based update rules. It is purely a topology dependent approach where the state of any node at time t, *S*_*i*_(*t*), is updated asynchronously based on the sum total number of activators and inhibitors incident on that node (Font-Clos et al. 2018). If the number of activators incident on the node is greater than the number of inhibitors, node is set to 1, otherwise 0. If the network state doesn’t change over time steps, it is said to have reached steady state. The evolution of network dynamics is governed by the following equation:

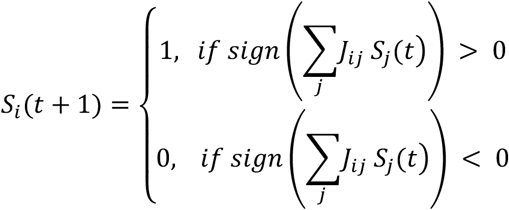

where, *j*_*ij*_, the edge from node *S*_*j*_ to node *S*_*i*_, is defined as,

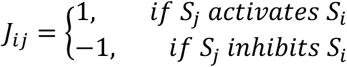

From both methods, we now have stable steady states as binary strings (0 → Low, 1 → High). We then characterize these steady states into hybrid or “terminal” (referring cumulatively to epithelial and mesenchymal states) based on an *EMT score* (Hari et al. 2021) (**Fig 1C**) and define the ‘hybridness’ of a network as fraction of steady state frequencies that can be classified as hybrid E/M (see details in materials and methods).

The results obtained from ODE based modeling show that the unperturbed EMP networks (referred to as “wild-type” or WT from hereafter) had low levels of hybridness as compared to most of their perturbed counterparts (**Fig 1D**). We quantified this comparison by calculating a percentile for WT hybridness in the distribution of perturbed hybridness values. We find that most EMP networks fall under the 25% mark (first quadrant) (**Fig 1E**). When simulated using the Boolean formalism, the hybridness of WT networks was near the median of the distribution for all networks (**Fig S2**). These results suggest that while WT EMP networks may have low hybridness intrinsically, minor (single-edge) perturbations in network-topology can altogether increase or decrease the hybridness in these networks. They also imply the role of network-topology in shaping the emergent dynamics, in particular hybridness, of many EMP networks. The relatively smaller frequency of hybrid states seen across all these networks can possibly explain a lower abundance of heterogeneous hybrid E/M phenotypes seen *in vitro* (George et al. 2017) and *in vivo* (Pastushenko et al. 2018).

### Change in global expression patterns correlates with change in hybridness

The observed changes in hybridness of WT networks upon single-edge perturbations triggered the investigation of characterizing the change in WT phenotypic distributions upon these perturbations. Thus, we analysed two metrics: *Jensen Shannon divergence (JSD)* and *J metric. JSD* quantifies the difference in the information contained in two probability/frequency distributions (Lin 1991). The *JSD* values lie in the closed interval [0, 1], with 0 indicating that the distributions are identical and 1 indicating that the distributions are altogether different (**Fig 2A**). While using *JSD* to compare the frequency distributions of phenotypes arising from WT networks with those arising from their perturbed counterparts, we identified specific perturbations that cause significantly higher changes in phenotypic distributions and hence have high *JSD* values (**Fig 2B**). For instance, when the inhibitory link from ZEB1 to GRHL2 is replaced by an activatory one (referred to as ZEB-GRHL2_2-1), the frequency of hybrid phenotypes increases from ∼10% to ∼45% (**Fig 2C, Table S1)**. On the other hand, when the inhibitory link from SNAIL to miR200 is replaced by an activatory one (referred to as SNAIL-miR200_2-1), there is minimal change in the phenotypic distribution for the perturbed network as compared to the WT one, thus the corresponding JSD is low (**Fig 2C, Table S1)**. An overall comparison of the frequency distributions of phenotypes for networks with high and low *JSD* with respect to the corresponding WT suggested that for EMP networks, the *JSD* upon perturbation is often reflective of an increase in hybridness of the network. A positive correlation between hybridness and *JSD* holds across different sizes of EMP networks (**Fig 2D**). In other words, change in the phenotypic distribution of EMP networks upon perturbation is associated with a significant increase in the frequency of hybrid phenotypes..

**Figure 2.**
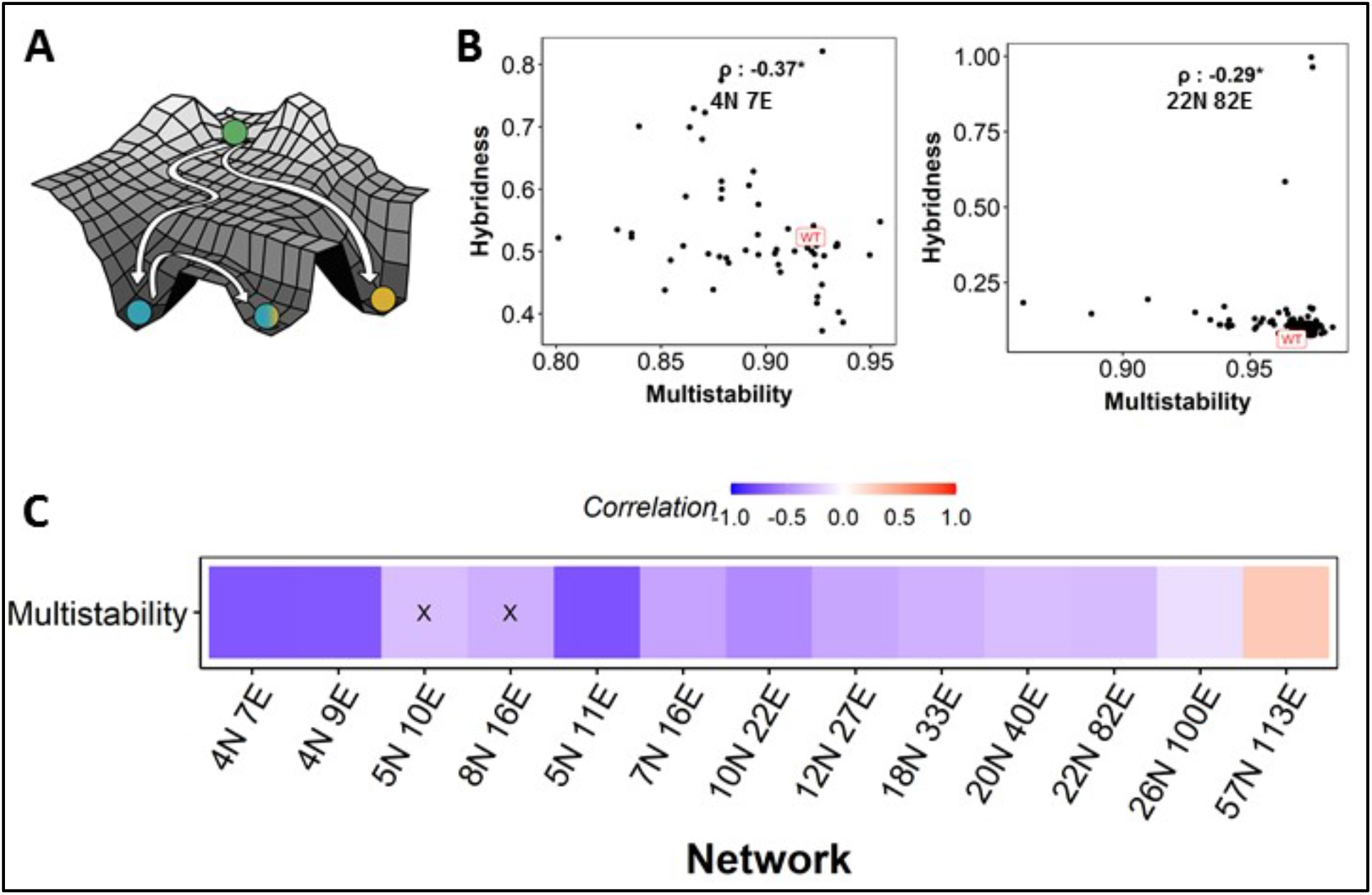
Increase in hybridness leads to reduced multistability. (A) Depiction of multistability, where valleys represent the steady states/phenotypes that the system (EMP network) can sample. **(B)** Representative scatter plots between multistability and hybridness. Spearman correlation coefficient is reported. **(C)** Heat map showing Spearman correlation coefficient values between hybridness and multistability across EMP networks. “X” represents p-value > 0.05.

The second measure, *J metric*, is a measure of cohesion in the expression levels between nodes of a network. We estimate *J metric* using the Pearson correlation matrix obtained by considering the steady state levels across all parameter sets (resp. the ensemble of initial conditions for Boolean). We take the absolute sum of all values in the upper triangle of the correlation matrix as the *J metric* of a given network (**Fig 2E**).

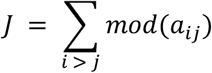

where *a*_*ij*_ is the any element of the correlation matrix *A* and represents correlation between nodes *i* and *j*.

In ODE based simulations, the WT EMP networks show high *J metric* as compared to their perturbed counterparts (**Fig 2F**). Given that the WT networks also show low hybridness, we probed whether these two metrics were correlated. For a given EMP network, we calculated *J metric* for all its corresponding perturbed networks as well as hybridness for them, and found *J metric* and hybridness to be negatively correlated (**Fig 2F, 2G**). This negative correlation is consistently seen considerably well in Boolean simulations **(Fig S3)**, endorsing that a characteristic feature of hybrid phenotypes is low cohesion between the node expression levels.

### Multistability increases with decrease in hybridness

Multistability is the ability of a system to achieve multiple steady states/phenotypes **(Fig 3A)**, thus enabling cells to reversibly switch to alternate phenotypes by adapting to environmental inputs. This ability to switch – also called as phenotypic plasticity - contributes to cancer aggressiveness and the success of metastasis (Gupta et al. 2019). It has been observed both *in vitro* as well as *in vivo* studies that hybrid phenotypes are less stable as compared to the “terminal” (epithelial and mesenchymal) phenotypes, i.e. they switch to other phenotypes more readily as compared to terminal phenotypes (Ruscetti et al. 2016; Pastushenko et al. 2018). In other words, hybrid phenotypes are proposed to be more “plastic” than the terminal phenotypes. Thus, we interrogated how the relationship between hybridness and multistability evolves upon perturbing EMP networks.

**Figure 3.**
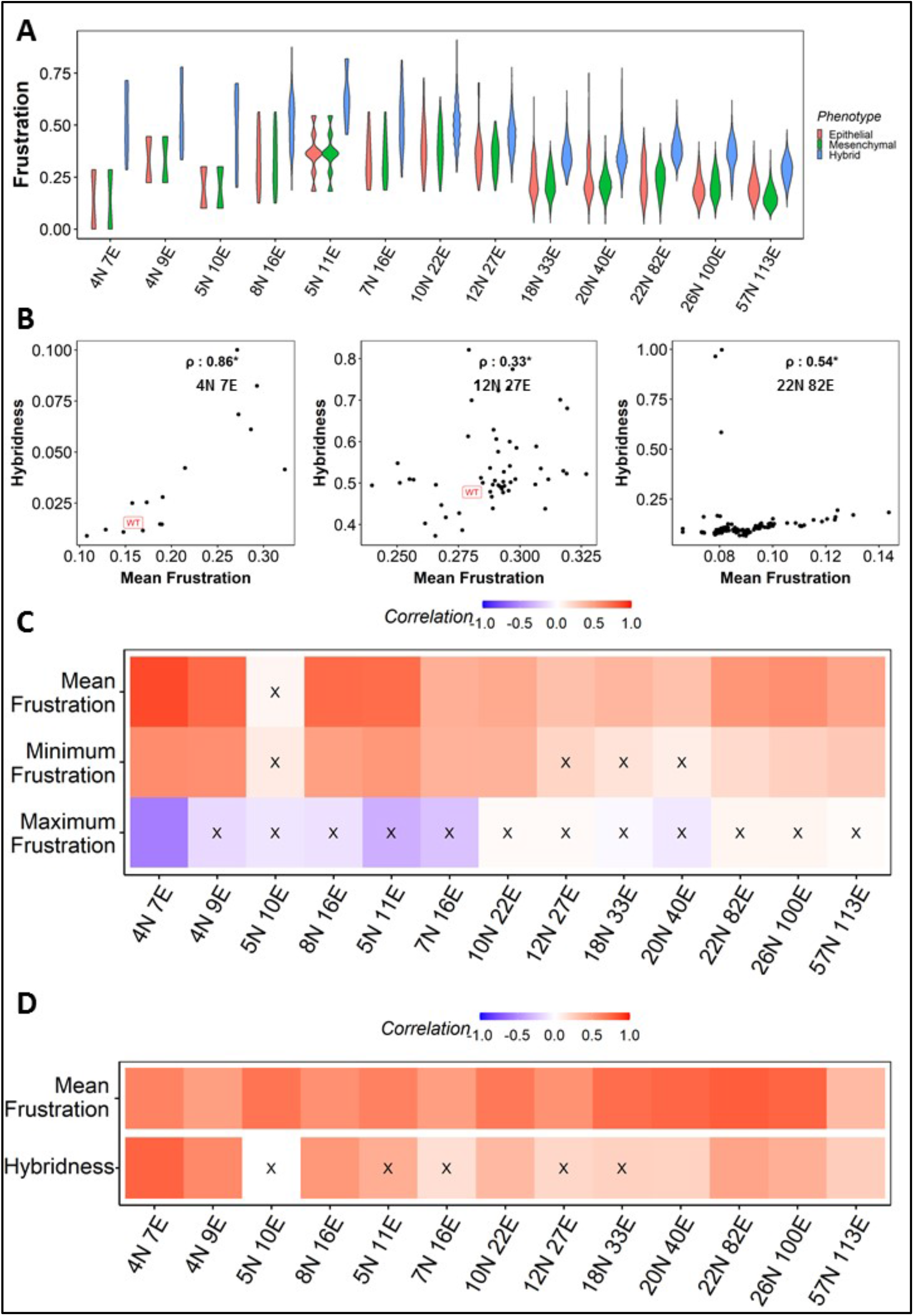
Frustration correlates positively with hybridness. **(A)** Violin plot depicting the distribution of frustration of Epithelial (peach), Mesenchymal (green) and Hybrid (blue) phenotypes (steady states) for WT EMP networks, showing that hybrid phenotypes have a higher frustration. **(B)** Representative scatter plots between Mean Frustration and Hybridness for (i) 4N 7E, (ii) 12N 27E and (iii) 22 N 82E EMP networks. Each dot is a perturbed network. Spearman correlation coefficient is reported. **(C**) Heat map depicting the spearman correlation of Hybridness with Mean Frustration (top row), Minimum Frustration (middle row) and maximum Frustration (bottom row). **(D)** Heat map depicting the spearman correlation of Predicted Frustration with Mean Frustration (top row) and hybridness (bottom row). “X”: p-value > 0.05

For RACIPE simulations, we estimate the multistability of a network by calculating the fraction of all the parameter sets that show multistability. For instance, if *P*_*MS*_ represents a parameter set that gives rise to multiple stable states and *P*_*T*_ represents the total number of parameter sets for which the network is simulated. We then define multistability (*M*) as:

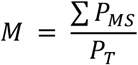

Our results show that WT networks are highly multistable compared to their perturbed counterparts (**Fig 3B**). This result is further confirmed by the negative correlation we found between hybridness and multistability across 12 of the 13 EMP networks **(Fig 3C)**. Overall, these results suggest that network topologies that support hybrid phenotypes might have a reduced capability of showing multistability.

### Hybrid phenotypes are highly frustrated

The tendency of hybrid states to be more “plastic” in wild-type network topologies can also be viewed as hybrid phenotypes being more “frustrated”, where frustration measures the extent of disagreement of a given steady state with the network topology (Tripathi et al. 2020b). Frustrated phenotypes can then be viewed as those resulting from gene expression patterns that may be the outcome of conflicting signals. Recent computational analysis of some EMP networks showed high frustration for hybrid phenotypes compared to the terminal phenotypes (Hari et al. 2021). We investigated whether the relationship between frustration and hybridness is maintained across perturbations in these EMP networks. Consistent with our previous observations, we noticed that the mean frustration of hybrid phenotypes was more than that for epithelial and mesenchymal ones, across the networks (**Fig 4A**). Here, the frustration of all steady states in all 13 EMP networks was calculated as follows. We first characterize an edge *j*_*ij*_ between nodes *S*_*i*_ and *S*_*j*_ to be frustrated (and denote it by *F*_*E*_) if

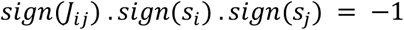

**Figure 4.**
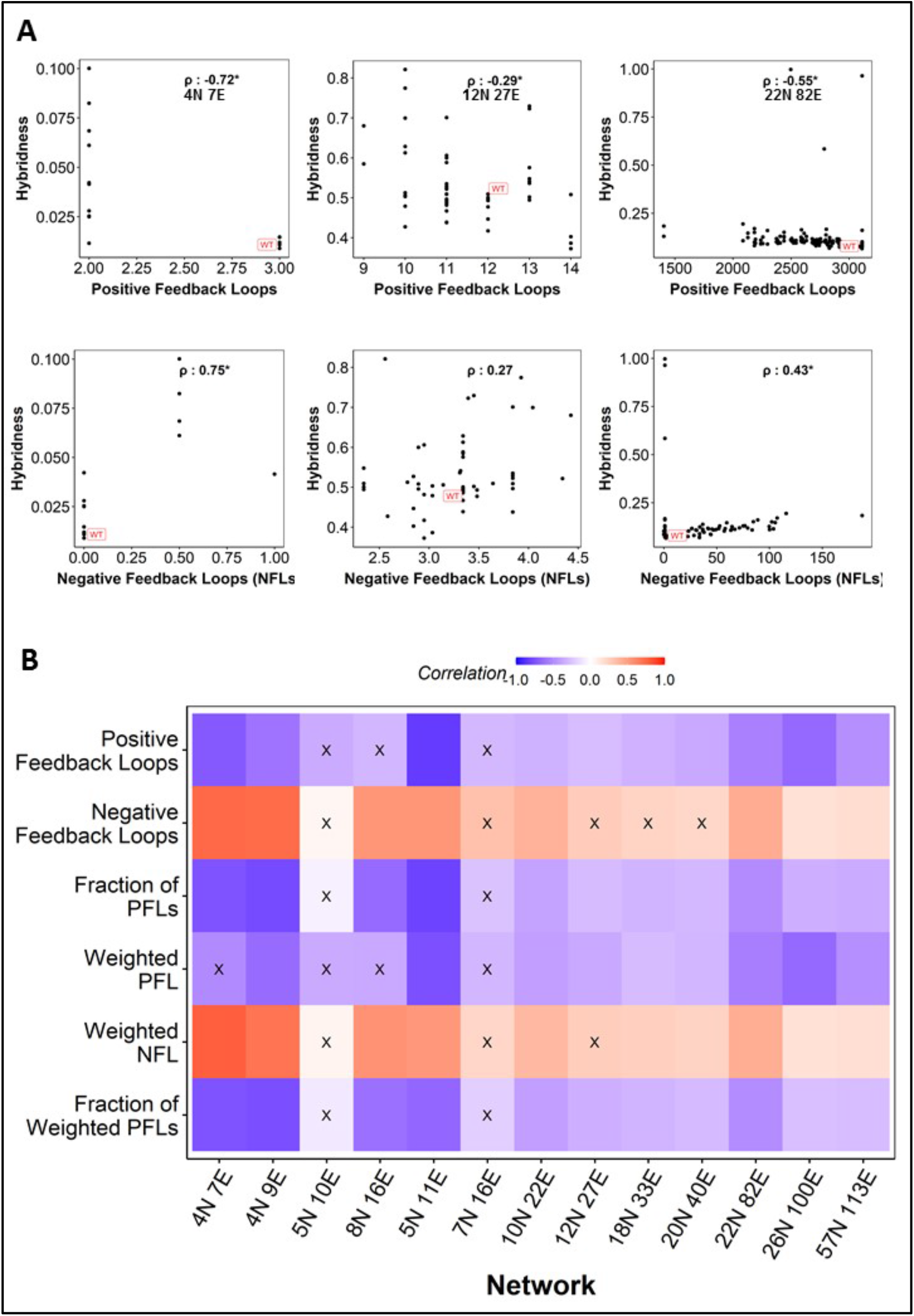
Positive feedback loops correlate with hybridness better than negative feedback loops. **(A)** Representative scatter plots between positive feedback loops (PFLs) and hybridness. Each dot is a perturbed network. Spearman correlation coefficient is reported. **(B)** Representative scatter plots between negative feedback loops (NFLs) and hybridness. Spearman correlation coefficient is reported. **(C)** Heat map depicting the spearman correlation between Hybridness and (from top to bottom) PFLs, NFLs, Fraction of PFLs, Weighted PFLs, Weighted NFLs and Fraction of weighted PFLs. “X”: p-value > 0.05

Here, *sign(j*_*ij*_) can be either *+1* or *-1*, depending on whether the edge is activatory or inhibitory. Similarly, sign of a node *S*_*i*_, *sign*(*S*_*i*_), can be either *+1* or *-1*, depending on whether the node is being activated or inhibited by the edges incident on node *S*_*i*_. We then define frustration of a state *S* (denoted by *F*_*S*_) as the fraction of edges that are frustrated in that state.

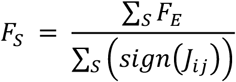

where, the numerator sums up the number of frustrated edges *F*_*E*_ in a state *S*, and denominator calculates the total number of edges involved in that state.

While obtaining frustrations of all the states, we then define frustration of a network *F*_*N*_ in three different ways, as the minimum, the maximum or the mean frustration of all states in that network.

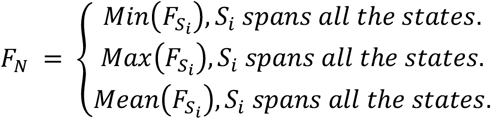

We calculated the minimum, maximum and mean frustrations for WT networks and their perturbed counterparts. Across the 13 EMP networks, we observe that while mean and minimum frustration correlated positively with hybridness, maximum frustration did not show any significant correlation **(Fig 4B, C)**. One possible way to interpret this trend is that the increase in hybridness upon perturbation can correspond to destabilization of states with least frustration, thus increasing the minimum and consequently the mean frustration value seen across the ensemble of states for a given network. The maximum frustration does not show strong trends in terms of correlating with hybridness, potentially of saturation in frustration values across the ensemble of steady states.

Frustration measures the disagreement between a steady state and the network topology (Tripathi et al. 2020b). We, thus, explored the possibility of identifying a network topology based metric to explain frustration. As there is a higher chance of frustration in a network having a higher number of negative feedback loops, we used the frequency of short-length (comprising less than and equal to 6 edges) negative feedback loops as a predictor of the frustration of a given network and called it as “predicted frustration” of a network. As expected, we saw consistently strong correlation between predicted frustration and the calculated mean frustration of a network, across all 13 EMP network topologies and their corresponding topological perturbations **(Fig 4D)**. However, the predicted frustration resulted in significant positive correlations with hybridness only in 8 of 13 EMP networks **(Fig 4D)**.

### Network topological metrics to explain hybridness

Given the observed association of predicted frustration with hybridness, we examined whether other network topology-based metrics, specifically involving feedback loops, correlate with network hybridness. The presence of positive feedback loops (PFLs) can enable multistability (Angeli et al. 2004) and we have demonstrated previously that an increase in the frequency of PFLs increases the likelihood of multistability emergent from a network (Angeli et al. 2004; Hari et al. 2020).

Across WT networks and their perturbed counterparts, for ODE based simulations, we found that the abundance of PFLs correlates negatively with hybridness **(Fig 5A, i-iii)**. This correlation was statistically significant across 10 of 13 EMP networks **(Fig 5B)**. Conversely, the total number of negative feedback loops (NFLs) were significantly positively correlated in 8 of the 13 EMP networks **(Fig 5A, iv-vi; Fig 5B)**. We next tested whether the fraction of PFLs (i.e. number of PFLs divided by the total number of NFLs and PFLs, for a given path length) correlates better with hybridness as a metric. It showed negative correlation with hybridness in 11 of 13 EMP networks, with the magnitude of correlation coefficient values equal to or more than PFLs for small and medium networks, as seen by the color intensity **(Fig 5B)**.

**Figure 5.**
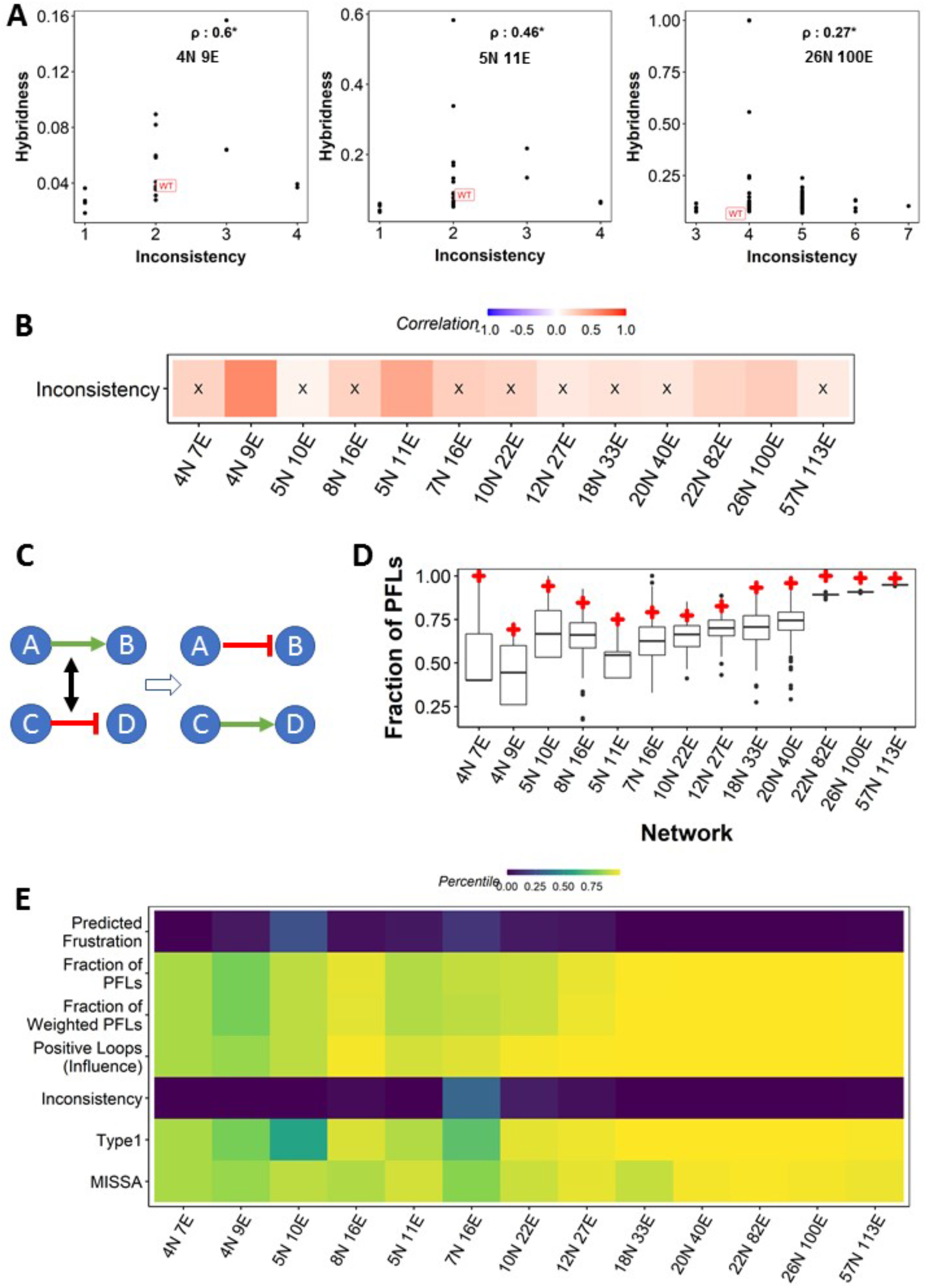
Inconsistency in EMP networks. (A) Representative scatter plots depicting a positive correlation between inconsistency and hybridness. (B) Correlation between inconsistency and hybridness obtained from ODE simulations of edge perturbations across EMP networks. (C) Random network generation schematic. (D) Distribution of inconsistency of random networks for all EMP networks. WT network inconsistency is labelled in red. (E) Heatmap showing the percentile position of WT networks in the distribution of random networks corresponding to the metrics considered in the current study. X: p-value > 0.05.

Given that the shorter loops are likely to be more effective, we took into account the importance of the length of the loops by assigning the loop-length (i.e., the number of edges in a loop) as a weight factor to the loop and calculated the number of weighted positive and negative feedback loops. We report that while weighted PFLs showed weaker correlation with hybridness as compared to their un-weighted counterpart, the weighted NFLs led to improved correlation as compared to un-weighted NFLs (10/13 significant) **(Fig 5B, S4)**.

### Higher order measures of feedback loops correlate better with hybridness

We next probed whether higher order measures of feedback loops can better explain hybridness. To calculate higher order feedback loops, we employed the influence matrix, a transformation of the interaction matrix that takes into accounts the influence of each node in the network on the other nodes, mediated through direct as well as indirect edges, with weights attached to the indirect influence based on the length of the corresponding paths (see materials and methods) (Chauhan et al. 2021). Thus, each cell in the influence matrix *I*_*ij*_ denotes how strongly node *i* “effectively” impacts node *j*, and has a value between -1 to +1, where -1 implies strong inhibition, +1 indicates strong activation, and 0 indicates no influence of node *i* on node *j*. We then identified positive and negative feedback loops between all pairs of nodes in the network using the influence matrix. Two nodes form a positive feedback loop if the influence between them in both directions is of the same sign, i.e. either they both effectively activate each other or inhibit each other (*I*_*ij*_ > 0 and *I*_*ji*_ > 0 or *I*_*ij*_ < 0 and *I*_*ji*_ < 0). However, if the influences are of opposite signs (i.e. *I*_*ji*_ > 0 and *I*_*ji*_ < 0 or *vice versa*), it is considered as a negative the loop. We define the strength of these loops as the product of influence values (*I*_*ij*_\**I*_*ji*_). A sum of these strengths is labelled as the positive loops (influence) or negative loops (influence) for a given network, as follows

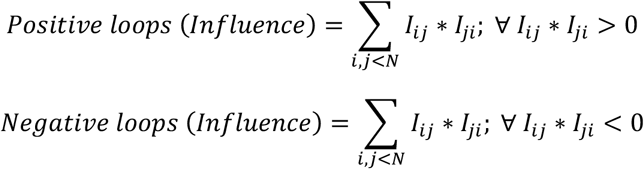

Where N is the number of nodes in the network.

The frequency of positive loops obtained from influence matrix showed negative correlation with hybridness (**Fig 6A**), significantly across 12 of the 13 networks analyzed (**Fig 6B**). The negative loops, however, did not show any such consistent patterns (**Fig 6B**).

**Figure 6.**
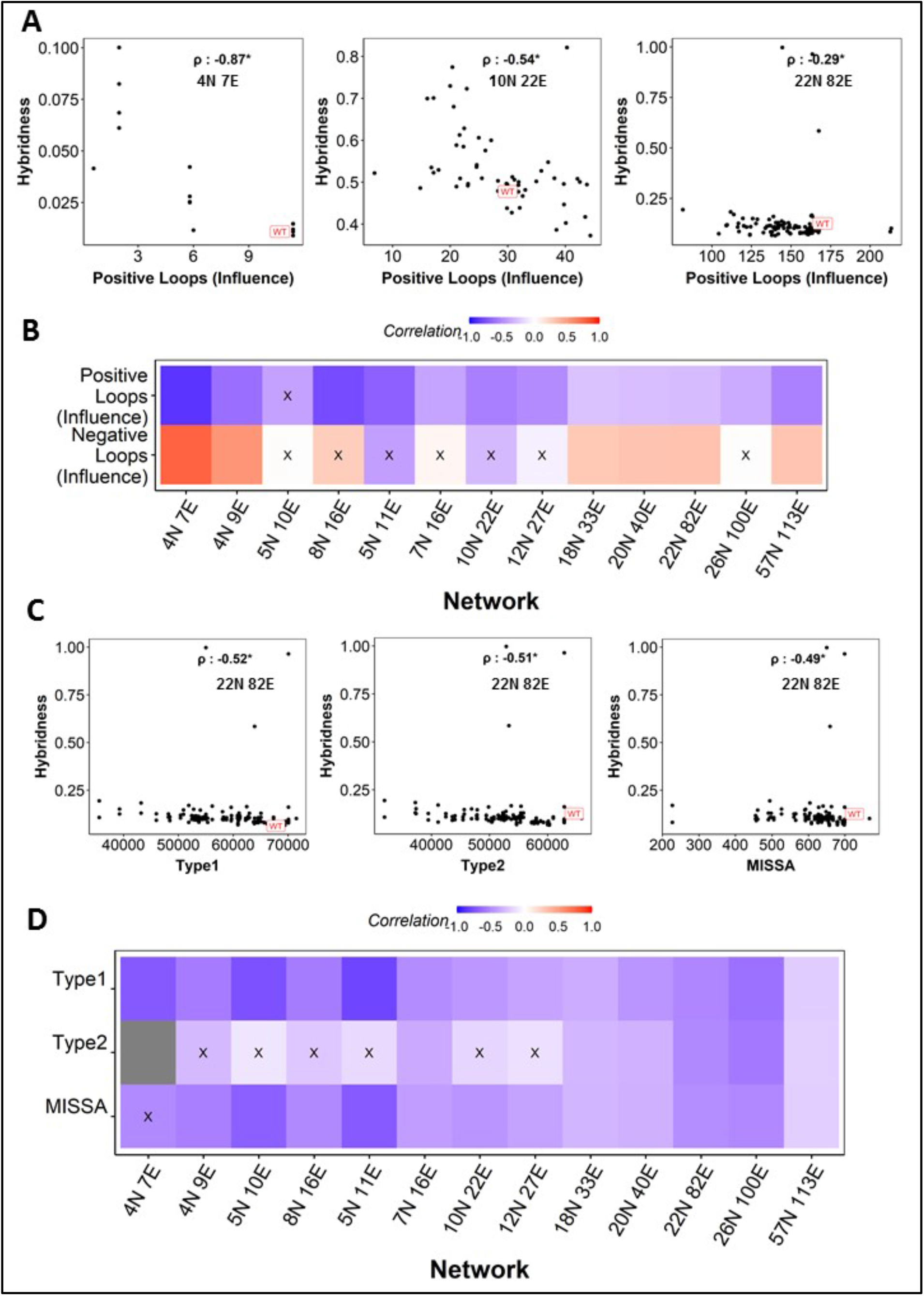
High-dimensional loop metrics perform better than simple loop metrics. **(A)** Representative scatter plots between positive feedback loops (PFLs) calculated from influence matrix and hybridness. Each dot is one perturbed network. Spearman correlation coefficient is reported. **(B)** Heat map depicting the spearman correlation between Hybridness and PFLs and NFLs calculated using Influence matrix. **(C)** Representative scatter plots between (i) Type-I, (ii) Tyep2 and (iii) MISSA HiLoop metrics and hybridness for 22N 82E network. Spearman correlation coefficient is reported. **(D)** Heat map depicting the spearman correlation between Hybridness and HiLoop metrics. “X”: p-value > 0.05

Further, we turned our attention to another form of higher order feedback loops – *HiLoops* (Nordick and Hong 2021), which are coupled feedback loops in diverse topologies. We chose three such metrics that consider purely PFLs: *Type-I*, where three PFLs are connected via the same node, *Type-II*, where three positive loops are connected via two nodes and MISSA, where two nodes form a double negative feedback loop, with one of the nodes having a positive feedback loop. Previous studies have shown that connected PFLs in EMT were capable of giving rise to (and sustaining) hybrid phenotypes (Jia et al. 2017b). Hence, we investigated if such motifs correlate with network hybridness. Our analysis revealed a negative correlation between the three *HiLoop* metrics and hybridness (**Fig 6C**). Among the three metrics, *Type-I* loops had strong negative correlation with hybridness, significant across all 13 EMP networks **(Fig 6D**). MISSA had a significant correlation in 12 networks and *Type-II* performed poorly, with correlations seen only in the larger networks (**Fig 6D**). These results indicate that coupled feedback loops can be used to get a better estimate of hybridness as opposed to individual feedback loops. Because the influence matrix takes into account paths of length more than one between a pair of nodes, the positive feedback loops obtained from the influence matrix can also be considered to be coupled, which could possibly explain more consistent trends obtained by influence matrix based loops compared to just PFLs (compare **Fig 6B** with **Fig 5B**).

These correlations obtained in ODE based simulations did not hold in Boolean simulations, however. While most of the networks showed insignificant correlation between hybridness and higher order loop metrics, *Type-I* did seem to have the best correlation patterns among all network topological metrics in Boolean simulations as well, leading to the conclusion that *Type-I* positive feedback loops are the best metric to understand trends in hybridness of EMP networks (**Fig S4**).

To check if stronger perturbations can reveal the trends between hybridness and the metrics studied, we perturbed the Boolean networks by randomly assigning weights to the edges of the EMP networks (i.e. its sign is maintained but the magnitude is randomly chosen between 0 and 1), similar to the parameter sampling in continuous-time simulations. We simulated the networks using the same mathematical formalism as before and obtained hybridness of each such randomly perturbed network. We determined the strength of feedback loops by multiplying the weights of the involved edges. We observe that the correlation between hybridness and mean frustration are stronger and more consistent than they were with single-edge perturbations (**Fig S5**). Since the topology does not change, the number of NFLs loops remain low in three of the networks, leading to the lack of correlation between predicted frustration and hybridness. The correlation with NFLs was more consistent, but the correlation with other metrics did not improve (**Fig S5**). Also, compared to small and medium networks, the correlations between *JSD* and hybridness in large EMP networks showed improved correlations thereby endorsing the trends seen for ODE based simulations.

### EMP networks are more consistent than random network counterparts

In the previous sections, we were able to use a metric based on NFLs to predict frustration. Furthermore, we find a positive correlation between such metrics and hybridness. In other words, the higher the network topology based frustration (as NFLs cause inconsistent connections between the pairs of nodes in a network), the higher the emergent hybridness.

Thus, we hypothesized that if perturbations were made to reduce the number of such inconsistent connections in an EMP network, it might be possible to reduce hybridness. To test this hypothesis, we calculated the “inconsistency” of a network, defined as the minimum number of edges whose sign must be flipped (activating edge turned to inhibition and vice-versa) to convert all inconsistent connections into consistent ones (Sontag 2007). Inconsistent connections include those involved in negative feedback loops and inconsistent feed-forward loops (estimated by counting the number of un-directed negative loops). Interestingly, the inconsistency in biologically observed EMP networks is found to be quite low (not more than 5 even in large networks with 100 edges) and correlates positively with hybridness **(Fig 7A)** across the networks **(Fig 7B)**. The lack of statistical significance of the correlation can be attributed to the small and discrete range of inconsistency in edge perturbed networks.

To investigate whether the network topological features that correlated with hybridness are unique to the network topology of WT EMP networks, we generated random networks by swapping edges randomly **(Fig 7C)** and compared the topological properties in these random networks with that of the WT network. We find that while positive-loop-based metrics showed a percentile of greater than 80 (i.e., more than 80% of random networks had a lower value of these metrics than that of WT **(Fig 7D)**, inconsistency and predicted frustration - metrics based on negative loops - in the networks showed a lower percentile **(Fig 7E)**. Together, these results suggest that WT EMP networks could have been designed to have lower fraction of hybrid states.

## Discussion

Reversible EMT is a dynamic cellular process underlying fundamental developmental programs which, however, goes aberrant in cancer cells to form metastasis. This cellular mechanism endows cancer cells the ability to undergo bidirectional switching between epithelial, one or more hybrid E/M, and mesenchymal cells. Promising computational and experimental studies have highlighted that hybrid E/M phenotypes exhibit relatively more plasticity than cells at the extreme ends, i.e., “pure” epithelial or mesenchymal cells, of EMT spectrum, thus abetting different steps of metastatic cascade (Biddle et al. 2016; Ruscetti et al. 2016; Font-Clos et al. 2018; Pastushenko et al. 2018; Biswas et al. 2019; Tripathi et al. 2020a). A better understanding of mechanism that enables them to enter into and exit from hybrid E/M cells would, thus, be a crucial step in improving therapeutic strategies targeting to curtail metastasis.

Recent studies while investigating the dynamics of gene regulatory networks underlying EMT have identified several PSFs that can stabilize one or more hybrid E/M phenotypes. A common lacuna of most of these studies is that they focused on influence of one particular gene on the emergence of hybrid E/M phenotype (Hong et al. 2015; Bocci et al. 2019; Silveira et al. 2020; Subbalakshmi et al. 2020, 2021; Norgard et al. 2021). Whether hybrid E/M phenotype is a consequence of the specific synergistic interactions of various genes in these networks, therefore, remains unidentified. Here we examined the underlying dynamics of 13 different GRNs identified for EMT/MET. Through a mathematical modeling approach, we found that all these networks are multi-stable and hybrid E/M phenotype is an innate feature of these networks, albeit at relatively lower frequency than the “pure” epithelial or mesenchymal phenotypes.

Making use of single-edge perturbations, we analysed the effect of various changes to network topology on hybridness. The WT EMP networks show a lower hybridness in comparison to the perturbed networks, as further confirmed by the correlations we obtained with *JSD* and *J metric*, measures of change in frequency distribution and cohesiveness respectively. A particularly interesting observation is the negative correlation between hybridness and multistability, counter-intuitive given that cells showing hybrid phenotypes have been shown to be highly plastic. This observation hints towards a possible trade-off between specialization provided by hybrid states (Cook and Wrana 2022) and bet-hedging opportunities provided by multistability. Such trade-offs are often seen in ecological systems (Egas et al. 2004) and have gathered interest in the context of heterogeneity in cancer cell populations too (Hausser et al. 2019). Given that the perturbations made to network topology (single-edge perturbations) are rather minor, these results can imply an evolution strategy for a cancerous population to survive by navigating through the specialist-generalist trade-off.

To identify the topological signatures (or design principles) that can explain association between the topology of GRNs and the hybrid E/M phenotype, we define different network-based metrics. Across these GRNs, we show that increasing the positive feedback loops increases the coherence among the network nodes that reduces the overall frustration within a network (or the states enabled by it). A completely opposite theme was observed in case of negative feedback loops - increasing the negative feedback loops reduces the coherence that in turn increases the overall frustration of the networks. Being highly frustrated, the probability that a hybrid E/M phenotype will occur from these networks increases significantly. Furthermore, we find that the PFLs and their combinations, as assessed by metrics *Type-I, Type-II, MISSA*, have a higher frequency in the WT EMP network topologies as compared to most of their randomized counterparts. Conversely, the NFL frequency and inconsistency, all of which measure the lack of coherence between the nodes of a network, are lower in WT EMP network topologies than most of their random network counterparts. Randomization can be interpreted as generating a sample of possible configurations in which the nodes of the network could have interacted over the course of evolution. The WT topologies then become the configuration that evolution seemingly has chosen over these random configurations. Extrapolating such interpretation, our results suggest that having coherent node interactions, and by extension, leading to a low fraction of hybrid states, may be evolutionarily preferred.

This analysis also holds significance in understanding the weak correlation between network topology metrics and hybridness obtained from Boolean simulations. RACIPE simulations show that hybridness correlates negatively with node-coherence metrics. Since WT networks are strongly coherent, the possibility of hybridness emerging from network topology itself (i.e., Boolean simulations) is quite low, making the correlation with topological metrics possibly weak. RACIPE, by sampling random parameters corresponding to different edge weights/node levels, can weaken the coherence and thus expose the correlation between topological metrics and hybridness better. Further support to this reasoning was provided by the improved correlations obtained in weight-based edge-perturbation Boolean simulations (in comparison to the edge-perturbation involving deletion/sign-change), because assigning random weights to the edges of the networks is an imitation of random parametrization employed in RACIPE.

Overall, our results emphasise that network-topology metrics alone can determine the phenotypic distribution emerging from these networks and presents an unorthodox mechanism – reducing negative feedback loops and increasing positive feedback loops - to restrict the emergence of hybrid E/M phenotype. These findings can provide first-hand insights to prevent different steps of metastatic cascade to curtail metastasis.

## Materials and Methods

### ODE based Model and Simulations

To investigate the underlying continuous-time dynamics of gene regulatory networks (GRNs), we used ordinary differential equation (ODE) based modeling approach implemented in *RACIPE* (Huang et al. 2017). *RACIPE* takes topology file of the GRN as input and simulates it in a continuous manner by constructing a system of ODEs. There is a one-to-one correspondence between the number of genes in the network and the number of ODEs generated by RACIPE. For instance, a simple network involving two nodes *X* and *Y* in a mutually inhibitory topology would be modelled by the following type of a system of ODEs.

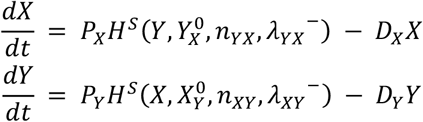

Here, *P*_*X*_, *P*_*Y*_ and *D*_*X*_, *D*_*Y*_ are the production and degradation rates of *X* and *Y*, respectively. Inhibition of gene *X* by gene *Y* is modeled as a shifted Hill function 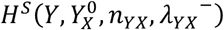 defined as:

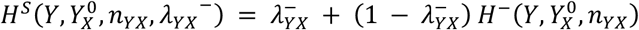

Here, 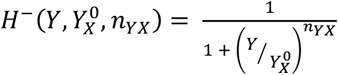 represents the inhibitory Hill function; *Y_X_*^0^ is the threshold level of *Y*; *n_YX_* represents the Hill coefficient of regulation; and 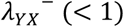 represents the fold change in the level of *X* caused due to the inhibitor *Y*. We can similarly model the inhibition of gene *Y* by gene *X*. The activation of *X* by *Y* is, however, represented by 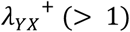.

The insufficient knowledge of kinetic parameters always poses questions on the validity of models. Here, that thing is taken care of by randomly sampling multiple parameter sets from predefined parameter ranges estimated from BioNumbers (http://www.bionumbers.hms.harvard.edu) (Milo et al. 2010), thereby generating ensemble mathematical models. Each ODE is solved numerically using *Euler* method of integration with step-size as *0*.*1*, number of initial conditions as *100*, and the number of iterations to solve ODE at each initial condition as *20*. These initial conditions are randomly sampled from a uniform distribution ranging between the lowest possible level and the highest possible level a gene can observe. To identify stable steady states, the cut-off for convergence of steady state is chosen to be *1*.*0*. For each network, we performed triplicate simulations using *10,000* parameter sets, thus generating *10,000* sets of ODE’s for each network, where each set has as many ODE’s as the number of nodes in the network. For instance, the smallest 4N 13E network will give rise to 10,000 sets of ODE’s where each set has 4 ODE’s. Likewise, the method follows in all the other networks.

### Normalizing and discretizing continuous-time ODE simulations

The output of mathematical models solved numerically through above simulation procedure are stable steady states. We used z-score standardization to transform the data such that the variables are fairly compared without any bias. The z-score of any stable state is estimated using below formula:

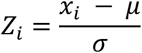

Here, *x_i_* is the *ith* stable state of any node; and, in that order, is the mean and standard deviation of all the stable states of that node.

Depending on whether the z-score of a steady state is positive or negative, these states are labelled as *1 (high)* and *0 (low)*, respectively. Any steady state is thus discretised and is represented by a string of *0’s* and *1’s*. For instance, steady states in a 3-node network would look like *“111”, “011”, “110”*, etc. However, if there are signal or input nodes - nodes on which no edge is incident from other nodes in the network - in a network, we don’t count those nodes in steady states. For instance, in a 3-node network with one node as signal node, a steady state would possibly be like *“11”, “01”, “10”, “00”*. Frequency of any steady state is calculated by counting its occurrence in all the parameter sets.

### Boolean Simulations

All Boolean simulations were carried out using the toolkit https://github.com/askhari139/Boolean.jl. Each network was simulated for 100000 initial conditions. All simulations were carried out in triplicates.

### Hybridness Calculations

Hybridness is a measure of the frequency with which a network gives rise to a hybrid phenotypic state. A hybrid state is defined as a cellular state in which one or more epithelial marker genes are simultaneously expressed with one or more mesenchymal marker genes. We estimate hybridness of a steady state in several steps: Firstly, we calculate correlation matrix of gene expressions of “wild-type” topology of a network. Secondly, the Epithelial and Mesenchymal genes in the network are identified through higher order clustering by using *hclust* function in *stats* package of *R 4*.*1*.*2*. Finally, we calculate Epithelial Mesenchymal Transition score (*EMT Score*) by subtracting average expressions of all the mesenchymal marker genes from the average expressions of all the epithelial marker genes, for each network.

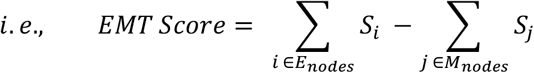

Here, *E*_*nodes*_ and *M*_*nodes*_ and in that order, denote epithelial and mesenchymal nodes in any network. Based on the *EMT score* that lies between *-1* and *1*, a state is characterized as hybrid if the *EMT score* lies between *-0*.*5* and *0*.*5*.

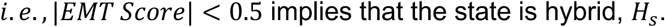

Hybridness of a network is then obtained by calculating the frequencies of the fraction of steady state frequencies that are hybrid. For instance, if *f*(*S_i_*) denotes the frequency of any stable steady state and *f*(*H_j_*) denotes the frequency of any hybrid state, we then define hybridness of a network as below:

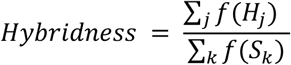

Where, *k* runs over all the steady states arising from a network and *j* runs over only those steady states that are hybrid.

Given that epithelial and mesenchymal marker genes in small networks (small number of nodes) are well established – miR200, OVOL2, GRHL2 are epithelial markers and ZEB, OCT4, KEAP1 are mesenchymal markers – we validated the above threshold for smaller networks by calculating their EMT scores based on the above markers. We found that the threshold was able to filter all the hybrid states while ignoring the “pure” epithelial and “pure” mesenchymal states.

### Single Edge Perturbations

In each of the 13 networks, we perturbed edges to create new networks – called as perturbed networks – by doing two types of edge perturbations: (1) We changed the sign of edges (one at a time). If the edge is activator, we changed it to a repressor edge and vice versa. (2) We delete any edge at a time to create new networks. Using this formalism, we will get 2*N perturbed networks from a network having N edges. To understand the role of topology in the network dynamics, all the perturbed networks were simulated using both ODE-based modeling approach as well as Boolean modeling approach.

### Comparing similarity between two frequency distributions using Jensen Shannon Divergence (*JSD*)

*JSD* is an information theory metric that quantifies the convergence or divergence of two distributions. For the two frequency distributions *f*_1_ and *f*_2_, *JSD* is mathematically defined as:

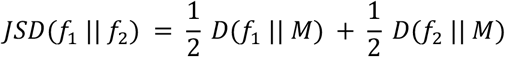

Where, 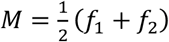 and the Kullback-Leibler divergence, *D*(*f*_1_ ‖ *f*_2_), is defined as below:

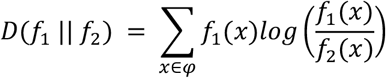

Here, *φ* is the probability space on which the probability distributions *f*_1_, *f*_2_ are defined. With *log*2 base, *JSD* values lie between *0* and *1*. Values close to *0* indicate overlapping distributions, while as values close to *1* indicate diverging distributions. All *JSD* calculations were performed by using *JSD* function implemented in *philentropy* package of *R 4*.*0*.*4*.

### Influence Matrix

Influence Matrix (*InfMat*) is a generalization of Interaction Matrix. While Interaction Matrix calculates the impact of any node in a network on all the other nodes that are directly connected to it, *InfMat* quantifies the influence of each node in a network on any other distant node through a predefined path length. Path length between two nodes is the number of edges through which these two nodes are serially connected. For instance, the influence matrix over path of length 2 can be defined as,

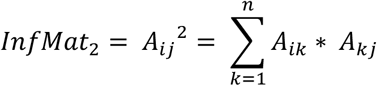

Here, the interaction matrix *A_ij_* considers influence of node *i* on node *j* through all paths of length 2. We accordingly generalize the case and define *InfMat* over path of length *l* as,

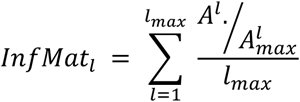

Here, 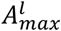– the normalizing factor – is the matrix by setting all non-zero elements of *A* to *1* and gives the maximum possible interaction between nodes in a network. It should be noted that the notation, “./”, is the division between corresponding elements of the matrices. Also, the summation is normalized by *l_max_* to restrict the range of the elements of *InfMat* between *-1* and *1*.

### Feedback Loops

In a directed network, a feedback loop is defined as a path formed of serially connected edges with the condition that the path originates and terminates at the same node. We say that a feedback loop is positive if it has zero or even number of negative edges (inhibitors); else, if a feedback loop has odd number of negative edges, the feedback loop is negative. We use *NetworkX* package in *Python3* to calculate positive and negative feedback loops in all the network considered in the study.

For feedback loops calculated using the influence matrix, we consider all pairs of nodes in a network. We then calculate the strength of feedback between the two nodes as the product of mutual influence values between the two nodes. If the strength is negative, we consider a negative feedback loop between the two nodes, and positive feedback loop otherwise. We then calculate the total number of such positive and negative feedback loops as follows:

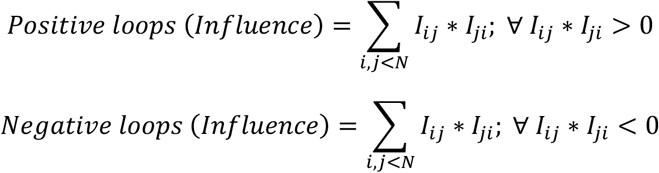

Where N is the number of nodes in the network.

### Higher Order Feedback Loops *(HiLoops)*

*HiLoops* are interconnections of two or more feedback loops. HiLoop toolkit (https://github.com/BenNordick/HiLoop) (Nordick and Hong 2021) was used to estimate three types of *HiLoops: Type-I*, where three positive feedback loops are connected through a common node, *Type-II, w*here a positive feedback loop between two nodes is such that each of these nodes is also a part of other disjoint positive feedback loops, and *Mutual Inhibition Single Self Activation (MISSA)*, where in a network topology the two nodes are mutually inhibiting each other and also one of these nodes is activating itself.

### Inconsistency

Using *NetworkX* package in python, we calculate the number of undirected positive and negative loops in a network. Then, we find out the common edges across the identified loops and iterate through the ones common between negative loops in the order of the number of loops they are common in and flip them one-by-one until all negative loops turn positive. The number of flips that are required to achieve this is labelled as inconsistency of a network. The code for calculation of inconsistency is attached in the github repository: https://github.com/csbBSSE/Hybridness-in-EMP-Networks.

## Code Availability

All codes are available at https://github.com/csbBSSE/Hybridness-in-EMP-Networks

## Acknowledgements

Mubasher Rashid is supported by SERB-NPDF *(PDF/2020/001235)* awarded by Science and Engineering Research Board (SERB), Govt. of India and by INSPIRE Faculty Fellowship *(DST/INSPIRE/04/2020/001492)* awarded by Department of Science and Technology (DST), Govt. of India. Kishore Hari is supported by Prime Ministers Research Fellowship, Govt. of India. Mohit Kumar Jolly is supported by Ramanujan Fellowship *(SB/S2/RJN-049/2018)*, SERB, Govt. of India and by the InfoSys Foundation, Bangalore.

## Author contributions

M. R., M. K. J. designed the research; M. R., K. H. performed the research; J. T. and N.K.S. contributed to data analysis; M. R., K. H., M. K. J. wrote the manuscript and discussed the results; all authors participated in revising the manuscript.

