## Supplementary Figures for "Network Topology Metrics Explaining Enrichment of Hybrid Epithelial Mesenchymal Phenotypes in Metastasis"

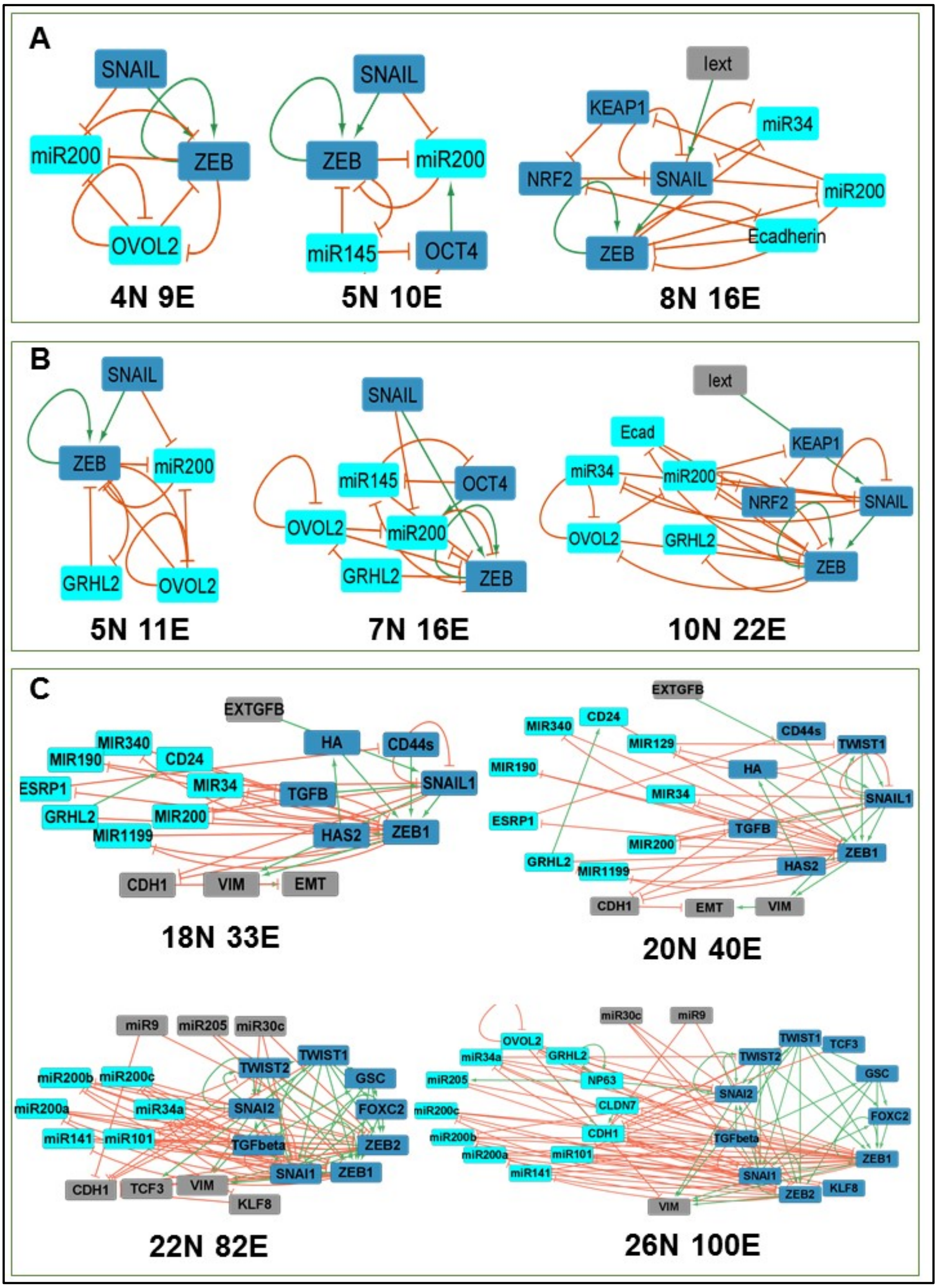


**Figure S1. EMP network topologies used for the analysis**. (A) Small size networks. (B) Medium size networks. (C) Large networks. Cyan colored nodes represent epithelial marker genes and blue colored nodes represent mesenchymal marker genes. Red edges represent inhibitions and green edges represent activations.

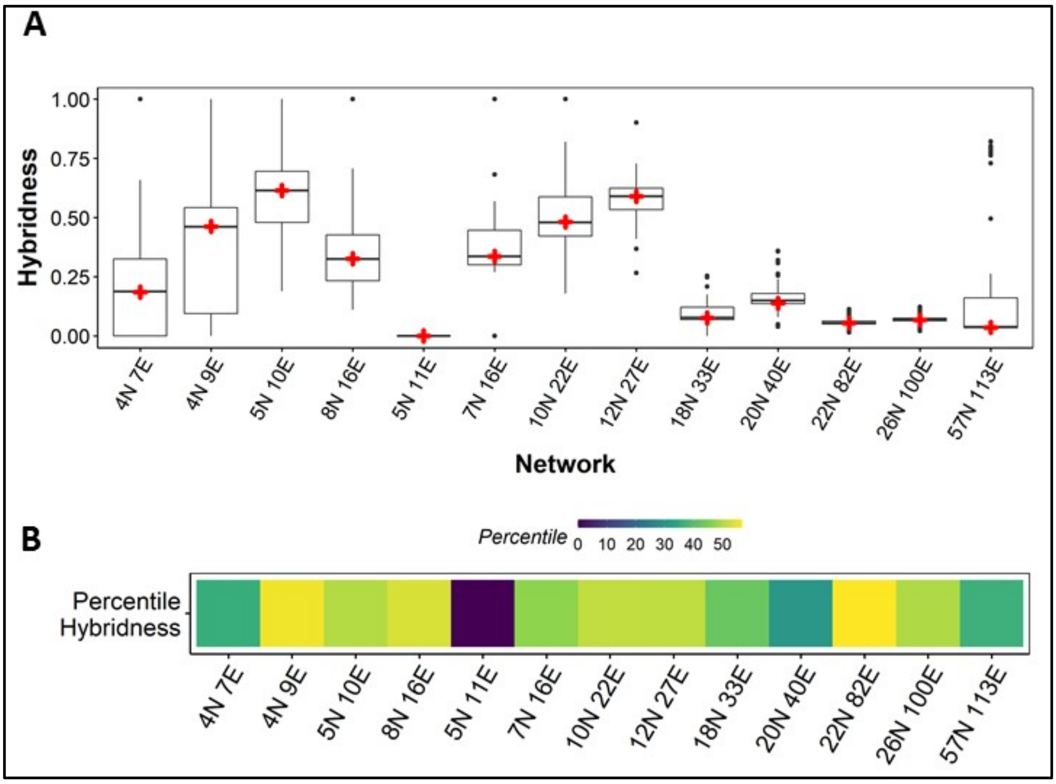


**Figure S2. Distributions of hybridness obtained from Boolean simulations**. (A) Each network is perturbed $2 * E$ times, where $E$ represents the number of edges, and hybridness of each perturbed network is calculated. Red mark represents hybridness of a WT topology of any network and the associated distribution is the hybridness of all of its perturbations. (B) Percentile hybridness of WT EMP networks in the distributions in S2A.

**Table S1. Edge perturbations and their interpretation for 4N 7E network.**

| ***Perturbation Index*** | ***Perturbation*** | ***Interpretation*** |
|---|---|---|
| 1 | miR200-ZEB_2-0 | Deleting inhibition of ZEB by miR200 |
| 2 | ZEB-GRHL2_2-1 | Changing inhibition of GRHL2 by ZEB to activation |
| 3 | ZEB-miR200_2-1 | Changing inhibition of miR200 by ZEB to activation |
| 4 | ZEB-GRHL2_2-0 | Deleting inhibition of GRHL2 by ZEB |
| 5 | ZEB-miR200_2-0 | Deleting inhibition of miR200 by ZEB |
| 6 | GRHL2-ZEB_2-1 | Changing inhibition of ZEB by GRHL2 to activation |
| 7 | miR200-ZEB_2-1 | Changing inhibition of ZEB by miR200 to activation |
| 8 | ZEB-ZEB_1-2 | Changing activation of ZEB by ZEB (self-activation) to inhibition |
| 9 | GRHL2-ZEB_2-0 | Deleting inhibition of ZEB by GRHL2 |
| 10 | ZEB-ZEB_1-0 | Deleting activation of ZEB by ZEB (self-activation) |
| 11 | SNAIL-miR200_2-0 | Deleting inhibition of ZEB by miR200 |
| 12 | SNAIL-ZEB_1-2 | Changing inhibition of ZEB by SNAIL to activation |
| 13 | SNAIL-miR200_2-1 | Changing inhibition of miR200 by SNAIL to activation |
| 14 | SNAIL-ZEB_1-0 | Deleting inhibition of ZEB by miR200 |

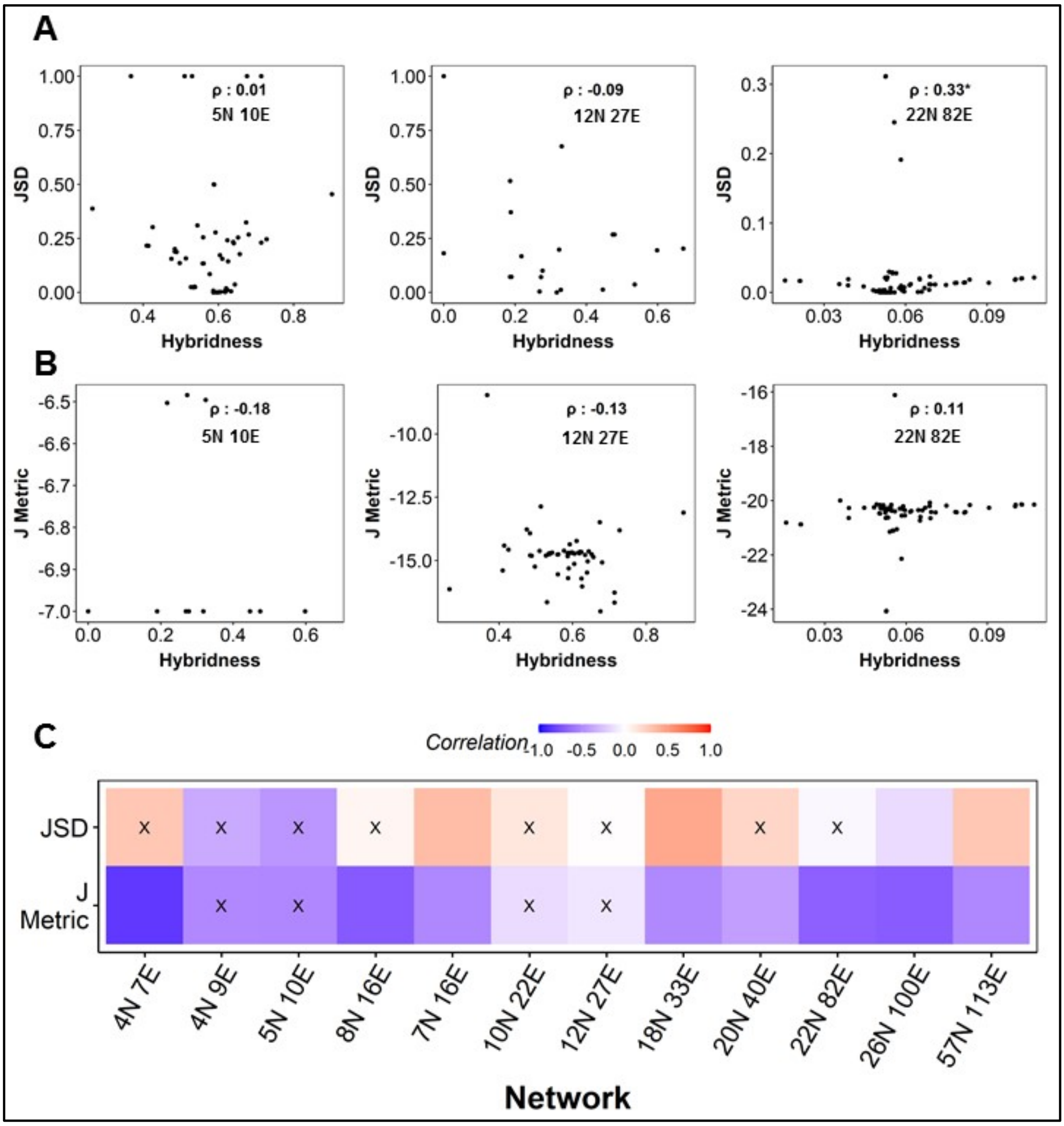


**Figure S3. Correlations of hybridness with *JSD* and *J metric* in Boolean Simulations.** (A) Representative scatter plots between *JSD* and hybridness. (B) Representative scatter plots between *J metric* and hybridness. (C) Spearman correlation between hybridness and *JSD* (top row) and *J metric* (bottom row) across the networks. "X" represents p-value > 0.05.

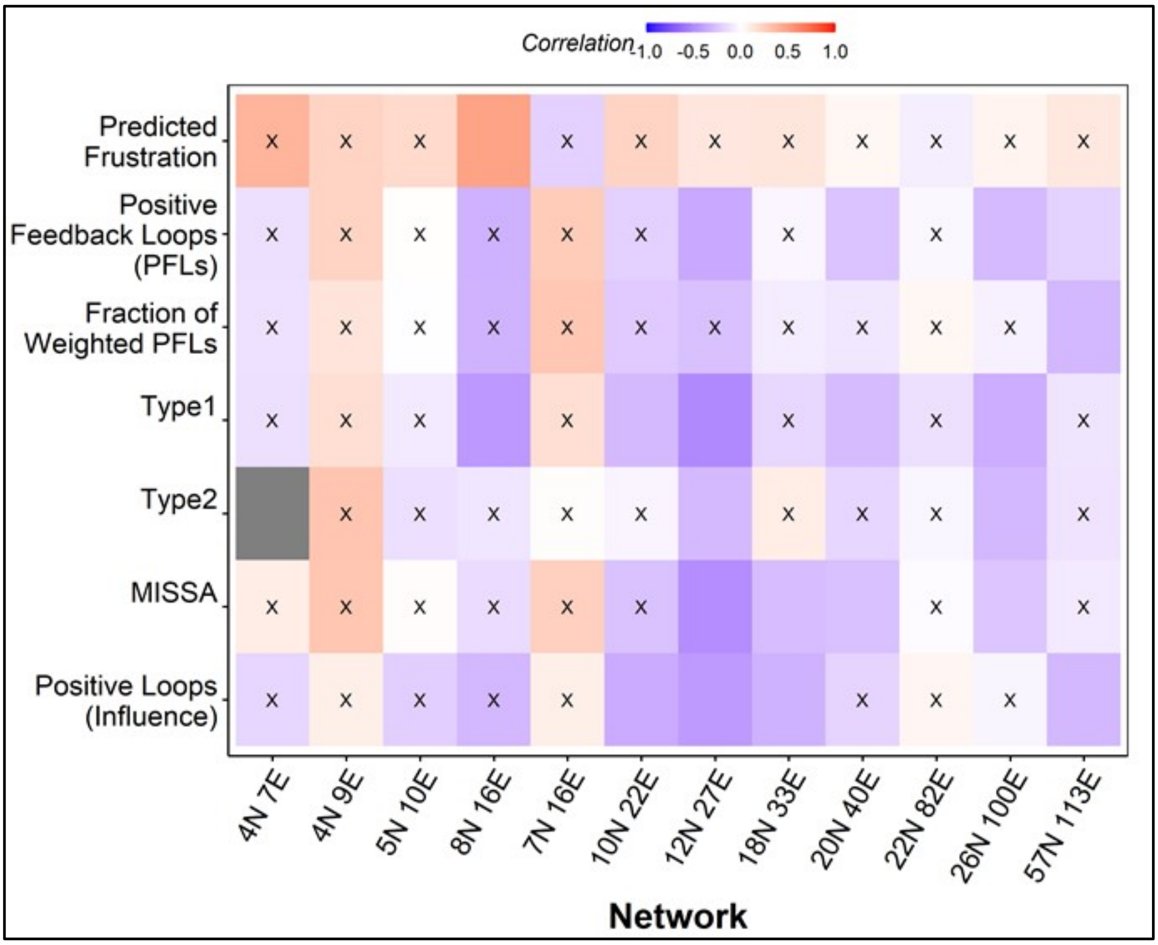


**Figure S4. Spearman correlation between Hybridness and loop metrics calculated from Boolean simulations**.

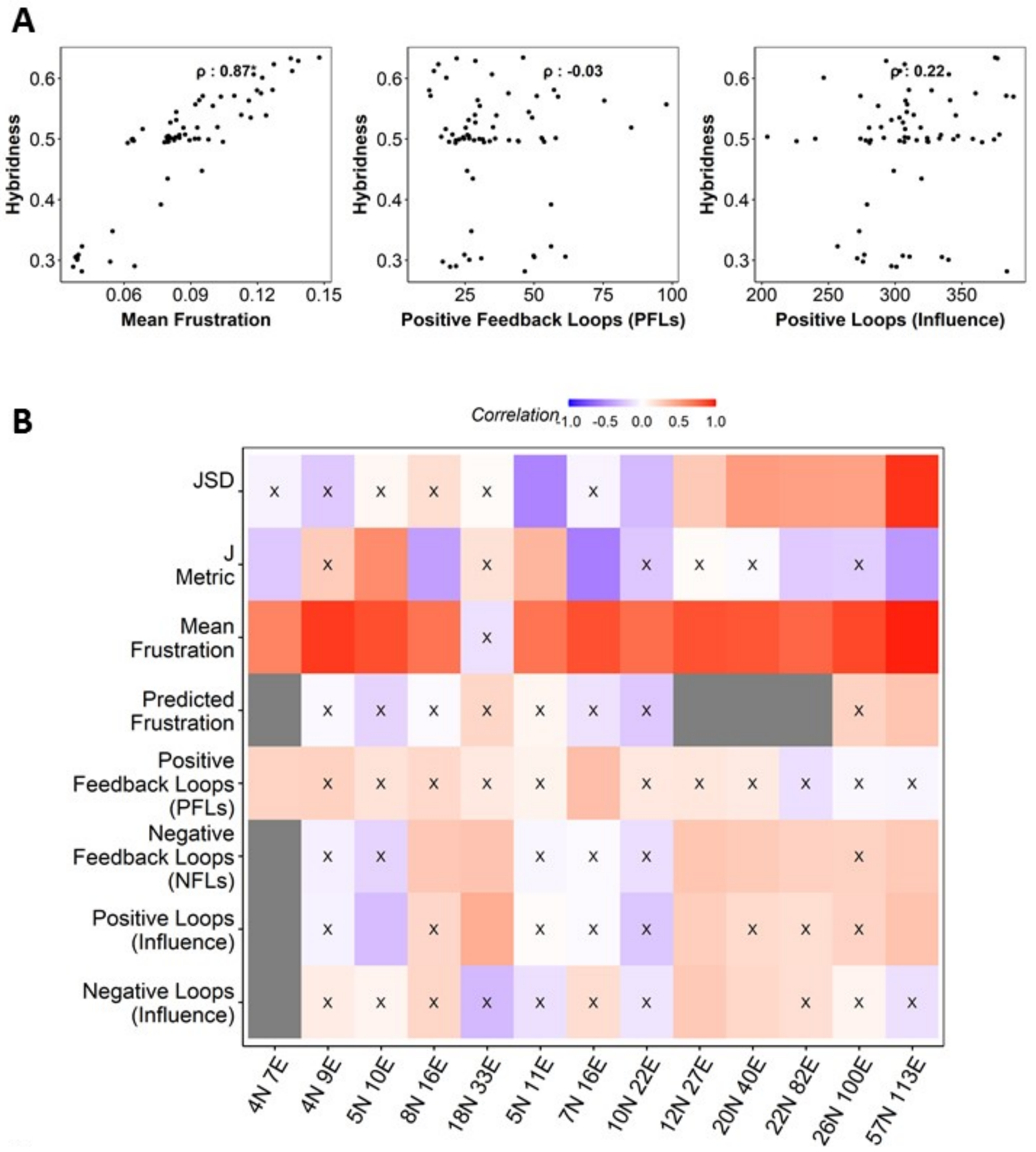


**Figure S5. Spearman correlation of Hybridness with frustration and loop metrics calculated from edge weight perturbations using Boolean simulations.**
